# Transcriptional profiling reveals T cells cluster around neurons injected with *Toxoplasma gondii* proteins

**DOI:** 10.1101/2020.06.09.143099

**Authors:** Emily F. Merritt, Hannah J. Johnson, Z. Sheen Wong, Adam S. Buntzman, Austin C. Conklin, Carla M. Cabral, Casey E. Romanoski, Jon P. Boyle, Anita A. Koshy

## Abstract

*Toxoplasma gondii*’s tropism for and persistence in the CNS underlies the symptomatic disease *Toxoplasma* causes in humans. Our recent work has shown that neurons are the primary CNS cell with which *Toxoplasma* interacts and infects *in vivo*. This predilection for neurons suggests that *Toxoplasma*’s persistence in the CNS depends specifically upon parasite manipulation of the host neurons. Yet, most work on *Toxoplasma*-host cell interactions has been done *in vitro* and in non-neuronal cells. We address this gap by utilizing our *Toxoplasma*-Cre system that allows permanent marking and tracking of neurons injected with parasite effector proteins *in vivo*. Using laser capture microdissection (LCM) and RNA-seq, we isolated and transcriptionally profiled *Toxoplasma*-injected neurons (TINs), Bystander neurons (nearby non-*Toxoplasma* injected neurons), and neurons from uninfected mice (controls). These profiles show that TINs transcriptomes significantly differ from the transcriptomes of Bystander and control neurons and that much of this difference is driven by increased levels of transcripts from immune cells, especially CD8^+^ T cells and monocytes. These data suggest that when we used LCM to isolate neurons from infected mice, we also picked up fragments of CD8^+^ T cells and monocytes clustering in extreme proximity around TINs and, to a lesser extent, Bystander neurons. In addition, we found that *Toxoplasma* transcripts were primarily found in the TINs transcriptome, not in the Bystander transcriptome. Collectively, these data suggest that, contrary to common perception, neurons that directly interact with or harbor parasites can be recognized by CD8^+^ T cells.

## Introduction

Obligate intracellular pathogens are dependent upon host cells for survival. Successful intracellular microbes, therefore, have highly evolved mechanisms to capitalize on host cell resources, avoid clearance by host cell-intrinsic defense mechanisms, and elude the recognition of infected host cells^1,2^. Our understanding of these host cell-microbe interactions primarily comes from *in vitro* studies and/or immune cells, which are relatively easy to isolate^3–6^. While such studies form the foundation of our understanding of host-microbe interactions, they have several limitations. These studies most commonly compare infected cultures to uninfected cultures^7–9^, which means that some differences assigned to “infection” are likely secondary to paracrine effects on surrounding cells (e.g. interferons). In addition, *in vitro* studies cannot replicate the multi-faceted interactions that occur *in vivo*, meaning these studies will miss pathways triggered only during *in vivo* infections. Finally, these studies are often conducted in cells not normally encountered by the pathogen (e.g. fibroblasts), meaning they may miss pathways that are specific to a subset of highly specialized host cells.

Such concerns are highly relevant for a microbe such as *Toxoplasma gondii*, an obligate intracellular parasite that has a wide range of intermediate hosts including humans and rodents. In most intermediate hosts, *T. gondii* establishes a persistent, long-term infection in certain organs and cells^10–13^. In humans and rodents, the central nervous system is a major organ of persistence^14^. This persistence and neurotropism underlies the parasite’s ability to reactivate to cause devastating neurologic disease in people with acquired immune deficiencies (e.g. AIDs patients^15–17^ or bone marrow transplant patients^18,19(p)^). Our *in vivo* understanding of CNS toxoplasmosis primarily comes from the mouse model, in which *T. gondii* preferentially interacts with and persists in neurons^20–23^. Thus, neuron-*T. gondii* interactions likely govern CNS outcomes, including *T. gondii* persistence. Yet very little is known about the neuron-*T. gondii* interaction, especially *in vivo*. What we do know about host cell-*T. gondii* interactions— that *T. gondii* secretes many effector proteins into host cells prior to and after full host cell invasion and that these effector proteins can be polymorphic between *T. gondii* strains, leading to strain-specific host cell manipulations— comes primarily from *in vitro* studies in fibroblasts and immune cells^3,24–32^.

To address this gap in knowledge, we sought to transcriptionally profile two types of neurons from *T. gondii*-infected mice— those directly manipulated by *T. gondii* and neighboring neurons that had not been manipulated by *T. gondii* but were still within in the same cytokine environment (Bystander neurons)— as well as neurons from uninfected mice. We reasoned that such comparisons would allow us to distinguish neuron expression differences arising from direct parasite manipulation from those changes elicited by the general neuroinflammatory response to *T. gondii*. To accomplish this goal, we utilized our *T. gondii* Cre system. In this system, Cre reporter mice that express a green fluorescent protein only after Cre-mediated recombination^33^ are infected with parasites engineered to express a *T. gondii*:Cre fusion protein that is injected into host cells concomitantly with other early effector proteins and prior to full invasion^34^. Thus, *T. gondii*-injected neurons, or TINs, permanently express GFP, while Bystander neurons or neurons from uninfected Cre reporter mice do not, allowing us to use laser capture microdissection to isolate these different groups of neurons. Finally, to enable our ability to identify both universal and strain-specific neuron manipulations, we utilized mice infected with either of the canonical, persistent *T. gondii* strains. These strains— belonging to either the type II and type III lineages— are genetically distinct, express different polymorphs of some injected effector proteins^24,29,31,30,28,25,27^ and are known to drive distinct CNS inflammatory responses^32^. Thus, with this combination of tools, we sought to define *T. gondii*’s strain-specific and universal effects on neurons during an *in vivo* infection.

## Results

### Isolation of neurons using laser capture microdissection

To develop insights into how neurons are manipulated by *T. gondii in vivo*, we intraperitoneally inoculated Cre reporter mice^33^ with saline (control), II-Cre, or III-Cre parasites (**Fig. 1**). At 21 days post inoculation (dpi), brains were harvested and processed for laser capture microdissection, including a rapid staining of sections with anti-NeuN antibodies to definitively identify neurons. We then used LCM to isolate 200 cortical neurons (NeuN^+^) from each saline-inoculated mouse, as well as 200 cortical *T. gondii*-injected neurons (TINs) (GFP^+^, NeuN^+^) and 200 cortical Bystander neurons (GFP^−^, NeuN^+^ and within 80-120 μm of a TIN) from each infected mouse (**Fig. 1**, **Fig. S1**). Each 200-neuron group was pooled and used for RNA isolation/library generation. The 25 libraries (5 control, 5 TINs II-Cre, 5 Bystander II-Cre, 5 TINs III-Cre, and 5 Bystander III-Cre) were sequenced at a projected depth of 50 million reads. High quality reads then underwent pseudoalignment, followed by differential gene expression analysis^35,36^. Our analysis showed that the total number of mapped reads was consistent across samples, with values ranging between 40-56 million, except for one III-Cre TIN sample which had approximately 16 million reads (**Table S1 & Fig. S2**). This sample grouped with the other TINs in PCA analysis (**Fig. S2C**) and thus was retained through all the subsequent analyses. To validate our isolation technique, we compared the transcript levels of the GFP protein expressed by these mice (ZsGreen) in the saline, bystander, and TINs samples. As expected, TINs samples showed 42 and 12.5 fold more GFP transcripts than saline and Bystander samples, respectively. Next, to determine if we had primarily isolated neurons, we analyzed the expression of known neuron-specific genes compared to known astrocyte and oligodendrocyte-specific genes^37^, the other two common parenchymal CNS cell types. Consistent with isolation of neurons, saline and Bystander neurons showed a higher abundance of neuron-specific genes compared to astrocyte and oligodendrocyte markers (**Fig. 2**). Unexpectedly, compared to the saline and Bystander neurons, TINs had a decrease in neuron-specific transcripts and an increase in astrocyte- and oligodendrocyte-specific transcripts (**Fig. 2**).

**Figure 1.**
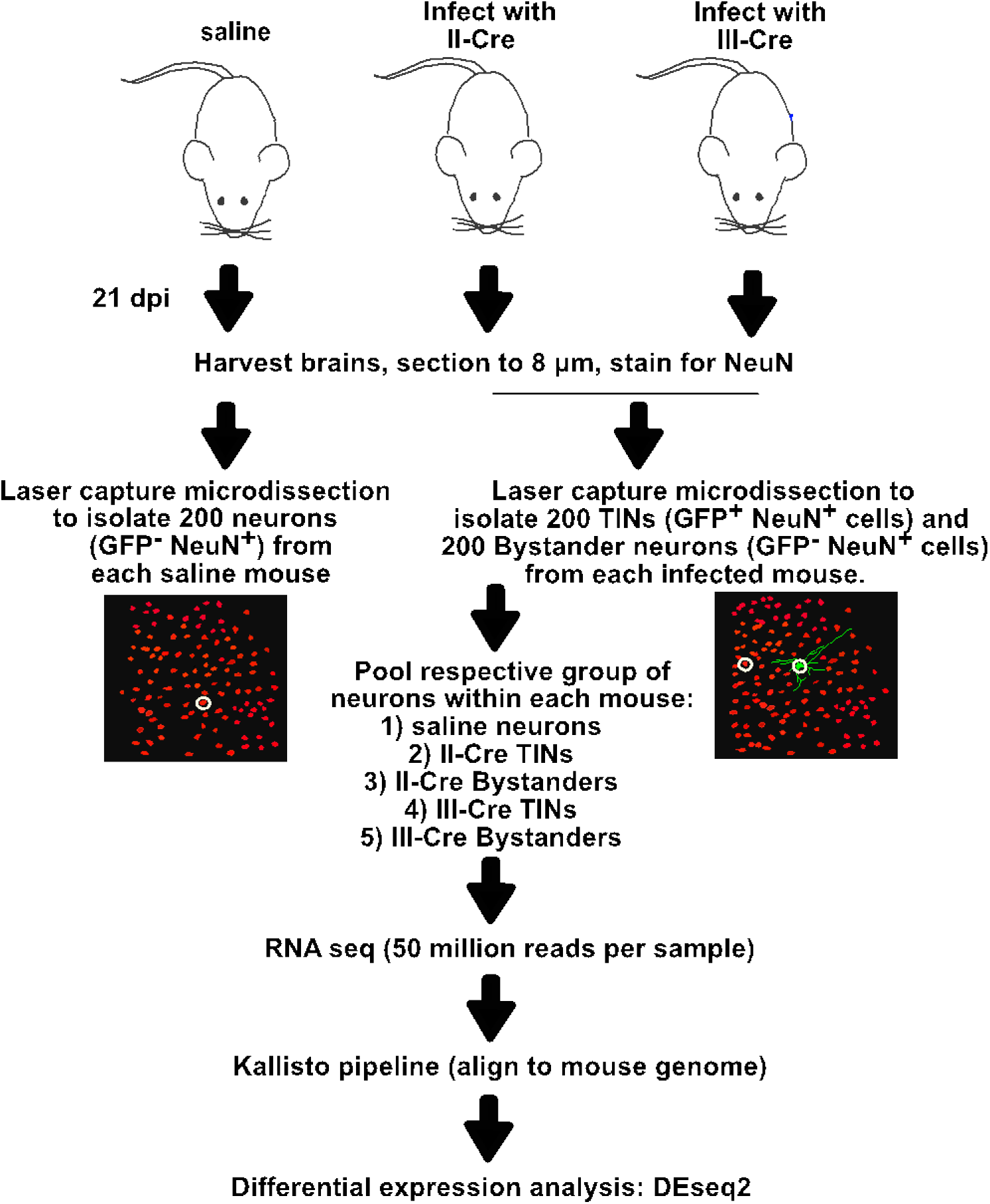
Schematic of Experimental Design. Mice were intraperitoneally inoculated with either saline or 10k syringe-released II-Cre, or III-Cre parasites. At 21 days post inoculation (dpi) brains were harvested, sectioned to 8 μm, and stained for neuronal marker, NeuN. LCM was used to isolate soma of 200 TINs and 200 bystander neurons from each mouse. 200 neurons were isolated from uninfected mice as a control. Each group of samples was pooled, sample quality assessed, and DNA amplified and aligned for RNA seq analysis.

**Figure 2.**
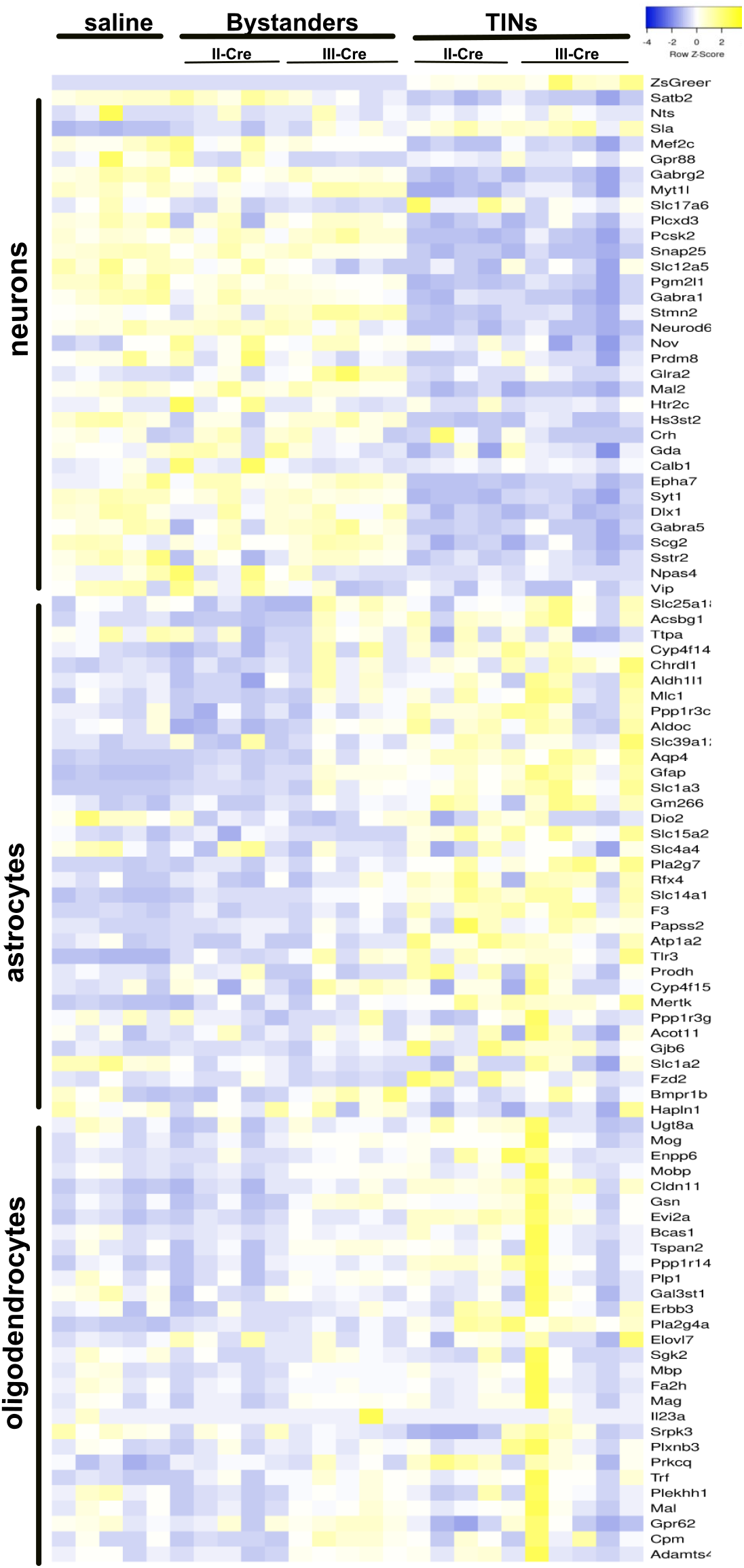
Neuronal transcripts are increased in saline and bystander samples, decreased in TINs. 5 samples for each condition, with 200 neurons per sample, were captured with LCM and sequenced with RNAseq. Raw reads were evaluated for known neuron, astrocyte and oligodendrocyte specific transcripts. Heatmap of cell-specific transcript reads; markers curated from Cahoy et al, 2008^37^.

Collectively, these findings show that 200 cells/sample generates high-quality transcriptional data. The high level of neuron transcript in saline neurons and Bystander neurons, in addition to the GFP in TINs transcriptomes suggest we isolated the cells we sought (cortical neurons, TINs, and Bystander neurons).

### Transcriptomes cluster by *T. gondii*-injection status not *T. gondii* strain status

As the decrease in neuron-specific transcripts in TINs did not segregate by infecting *T. gondii* strain type, we sought to determine if this segregation by group—but not parasite type—also held true at a global level. To investigate this possibility, we performed a Principal Component Analysis (**Fig. S2C**), using all genes with different transcript abundance. Consistent with our findings for parenchymal CNS cell transcripts, saline and Bystander samples clustered together, while TINs were the most distinct group and showed the most within group variability. Again, neither Bystander nor TINs samples segregated by infecting *T. gondii* strain (i.e. all TINs cluster together and all Bystanders cluster together), suggesting that global, common differences obscure smaller, strain-specific differences in our data.

We identified 7092 genes that differed in our infected groups from saline (**Table S2**), with marked differences between Bystanders and TINs (**Fig. 3A**). TINs showed a higher number of unique transcriptional changes (2081 upregulated, 2039 downregulated) compared to Bystanders (98 upregulated, 225 downregulated) (**Fig. 3B, C**, **Table S2**). Collectively, these findings are consistent with prior studies of infected non-neuronal cells^3,38,39^, indicating that injection with *T. gondii* effector proteins (and possible infection) caused dramatic alterations in the transcriptional landscape.

**Figure 3.**
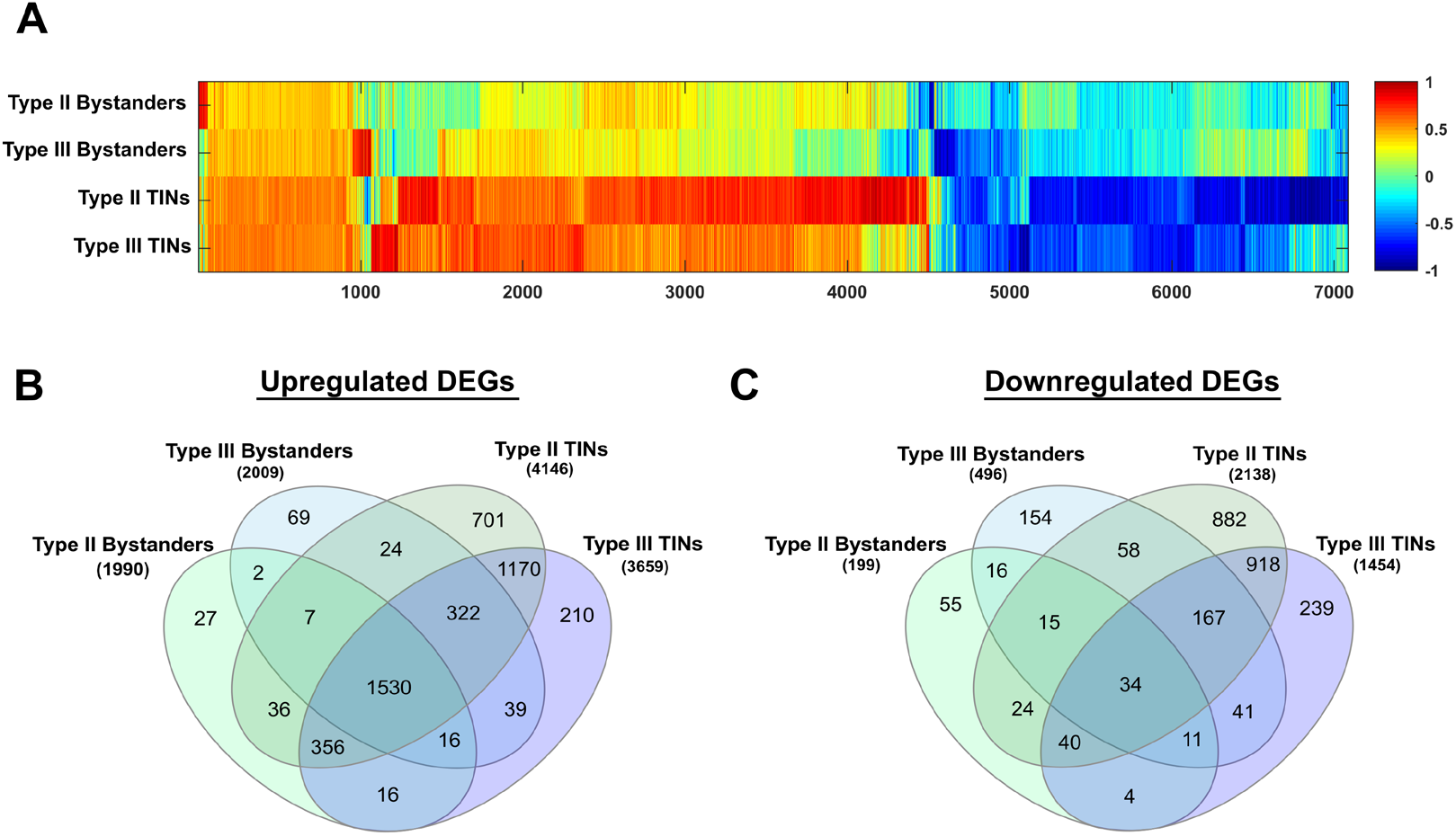
*Toxoplasma*-injected neurons (TINs) cluster together and have the highest number of differentially expressed genes. Comparison of 7092 genes ≥ 2-fold change and with a false discovery rate of 0.05 at padj < 0.05 across groups, compared to saline controls. **A)** Heat map of 7092 genes, Bystander and TIN groups normalized to saline with log_2_ fold changes, x-axis indicates number of genes. The log_2_ values of genes were clustered and normalized with the Euclidean norm with the default settings of MarVis suite. **B)** Upregulated DEGs. **C)** Downregulated DEGs.

### Pathway Analysis reveals enrichment for immune pathways in TINs versus Bystanders

To investigate the function of the DEGs in the Bystander neurons and the TINs, we used Ingenuity® Pathway Analysis (IPA®) to conduct a core analysis on each group and then ran a comparison analysis between groups, all normalized to saline. With no filtering, immune pathways dominated the list (**Table S3**). To delve into these differences, we filtered our IPA analysis settings to immune and inflammatory pathways. As expected, these pathways were markedly increased in all groups from infected mice compared to the control group (from saline-inoculated mice) (**Fig. 4A**, **Table S3**). In most pathways, TINs had higher levels of expression of the genes within a pathway, compared to Bystanders (**Fig. 4A**). These data suggested that both Bystander neurons and TINs were in a highly inflamed CNS environment with TINs showing relatively more markers of inflammation.

**Figure 4.**
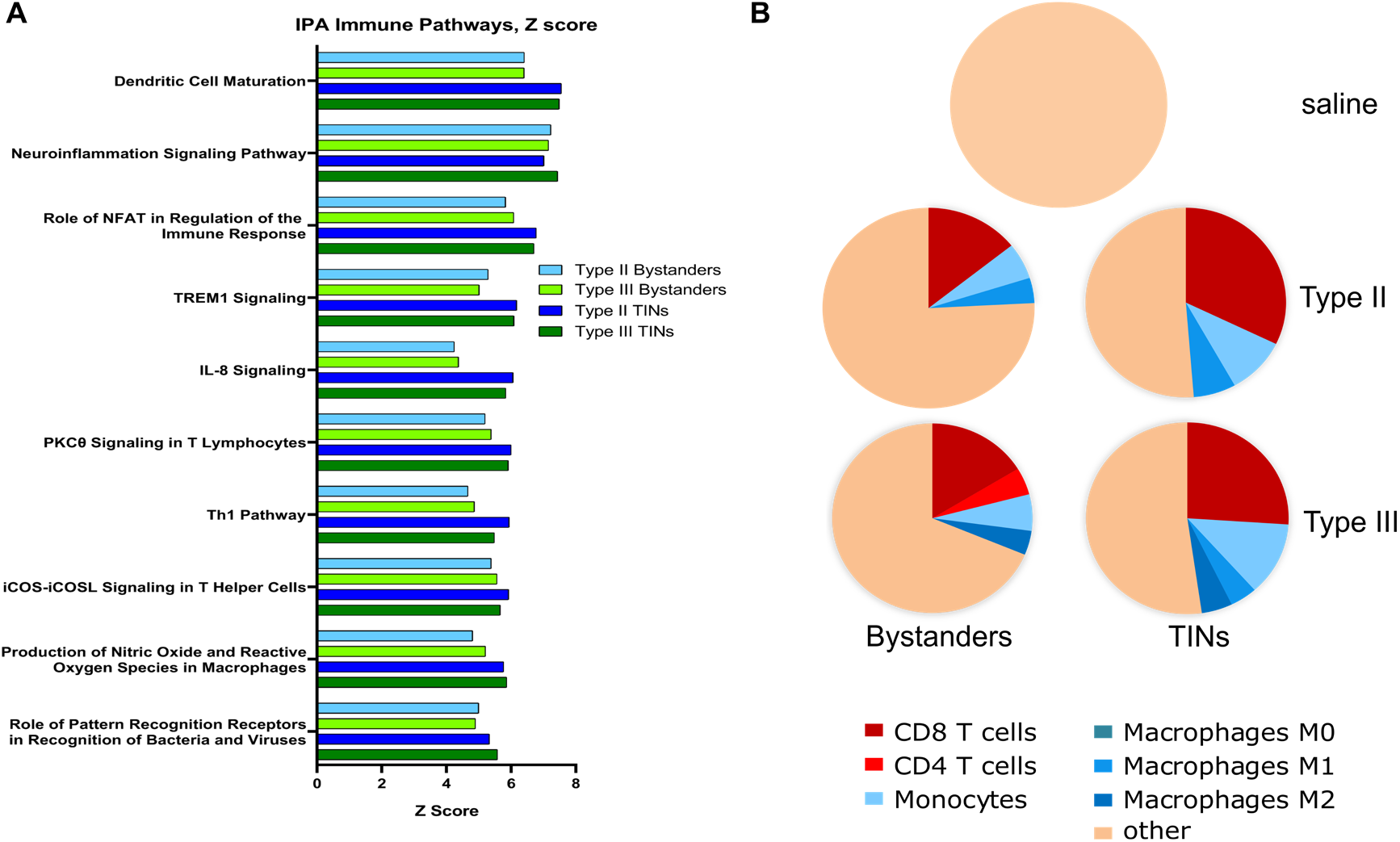
Immune cells cluster around TINs compared to Bystanders and saline. Z score of upregulation levels of all genes associated with pathways compared to saline. All pathways shown have p-values < 0.05. **A)** Top 10 immune pathways for Type II Bystanders, Type III Bystanders, Type II TINs and Type III TINs. **B)** Pie charts of CIBERSORT deconvolution using LM22 basis matrix. Each pie chart represents the average proportion of specific immune cell types in the identified samples. Only immune cell types with estimated abundance of ≥ 4% are shown. 4% was selected as the cut off because the average of the saline samples had a p-value of 1 and all immune cell abundance estimates in the saline samples were < 4% (**Table S4**). The p-values for the average of all infected deconvolutions was <0.05 (**Table S4**). Population denoted as “other” represents neurons, astrocytes, oligodendrocytes, microglia, and immune cell estimates that fell below 4%.

When we examined the up-regulated genes in the neuroinflammation pathway, we found many genes that are common in immune responses and produced by many cell types. Such genes include NFκB, STAT1, MHCI, TNF, IFN-β, potentially suggesting that neurons might be mounting typical cellular immune responses to infection and cytokine stimulation^7^ and that TINs, the cells with intimate *T. gondii* contact, received more cytokine signaling and thus generated more robust cytokine responses. But such a hypothesis did not explain the enrichment for immune cell-specific pathways. For example, the Th1 pathway has genes specific to Th1 cells (CD3, CD4), which would be highly unlikely to be expressed by neurons (even in the setting of inflammation). This pathway, as well the dendritic cell maturation pathway, suggested that when we used LCM to isolate neurons, we might have also picked up bits of immune cells clustering around Bystander neurons or TINs.

To formally assess this possibility, we analyzed our DEGs with CIBERSORT^40^, an algorithm that allows one to estimate the abundance of specified cell types based on gene expression. We performed the deconvolution using the LM22 signature matrix, which can deconvolve a wide range of immune cells, including different macrophage and T cell subsets (e.g. M1 macrophages). As expected, CIbersort analysis did not identify significant levels of immune cell transcripts in the saline group (p=1.0, **Table S4**). The lack of evidence for macrophage transcripts suggests that the saline transcriptomes have very little contamination with genes from microglia, the tissue resident macrophages of the CNS (**Fig. 4B**, **Table S4**). Conversely, the transcriptomes from infected mice— both Bystander and TINs— showed evidence for moderate-to-high levels of T cell-, monocyte-, and macrophage-specific transcripts. These data are consistent with prior evidence that, during CNS infection with *T. gondii*, these cell types infiltrate into the CNS and significantly contribute to the control of CNS toxoplasmosis^41,42,10,43–47^. Consistent with TINs showing higher levels of immune pathways (**Fig. 4A**) and lower levels of neuron transcripts compared to neurons from saline-inoculated mice or Bystander neurons (**Fig. 2**), the immune cell percentages were increased when comparing TINs to bystander neurons (**Fig. 4B**, **Table S4**).

Collectively, these data are consistent with previous work which has shown upregulated inflammatory pathways and an increase in immune cell infiltrate during infection^48–51^. Our identification of immune cell transcripts within our “neuron” transcriptomes suggest our samples contain transcripts from non-neuronal cells, such as CD8^+^ T cells, that were in extremely close proximity to the isolated neurons.

### Parasite transcripts are primarily found in TINs transcriptome

Given that our TINs transcriptome contained a higher number of transcripts from immune cells compared to our Bystanders, we sought to determine how parasite transcripts partitioned between TINs and Bystanders. After eliminating all reads that mapped to the host transcriptome (see Materials and Methods), we mapped the remaining reads against the *T. gondii* genome and quantified putative parasite transcript abundance in all samples (including neurons from saline-injected controls). We first eliminated all genes that had more than 20 total reads across all 5 saline-injected samples (as any significant mapping to the parasite transcriptome in these samples would represent genes with high similarity between parasite and host), leaving 8,605 transcripts that we analyzed in a number of ways. To qualitatively assess the quantity of parasite transcripts in TINs versus Bystanders, we further filtered the data to include only genes with at least 1 read in 15 of the samples leaving 527 genes, for which we calculated log_2_-transformed fragments per million reads mapped to the host transcriptome (**Table S5**). We did this to normalize each sample independent of the number of mapping parasite reads so that the samples could be compared independently of infection status and to use only genes of comparatively high abundance in the TINs. The TINs cell populations had consistently higher read counts for parasite transcripts compared to Bystander cells, with the exception of sample IIIT_1 (**Fig. 5A**). This finding was also reflected in the PCA analysis (**Fig. 5B**), where all TINs samples except IIIT_1 clustered together along the major PC1 axis, while all Bystander samples except IIIB_2 clustered along this same axis with saline-injected control samples (unlabeled cluster on left; **Fig. 5B**). These data show that, as expected, TINs harbor the majority of parasite transcript sequenced from LCM-captured cells.

**Figure 5.**
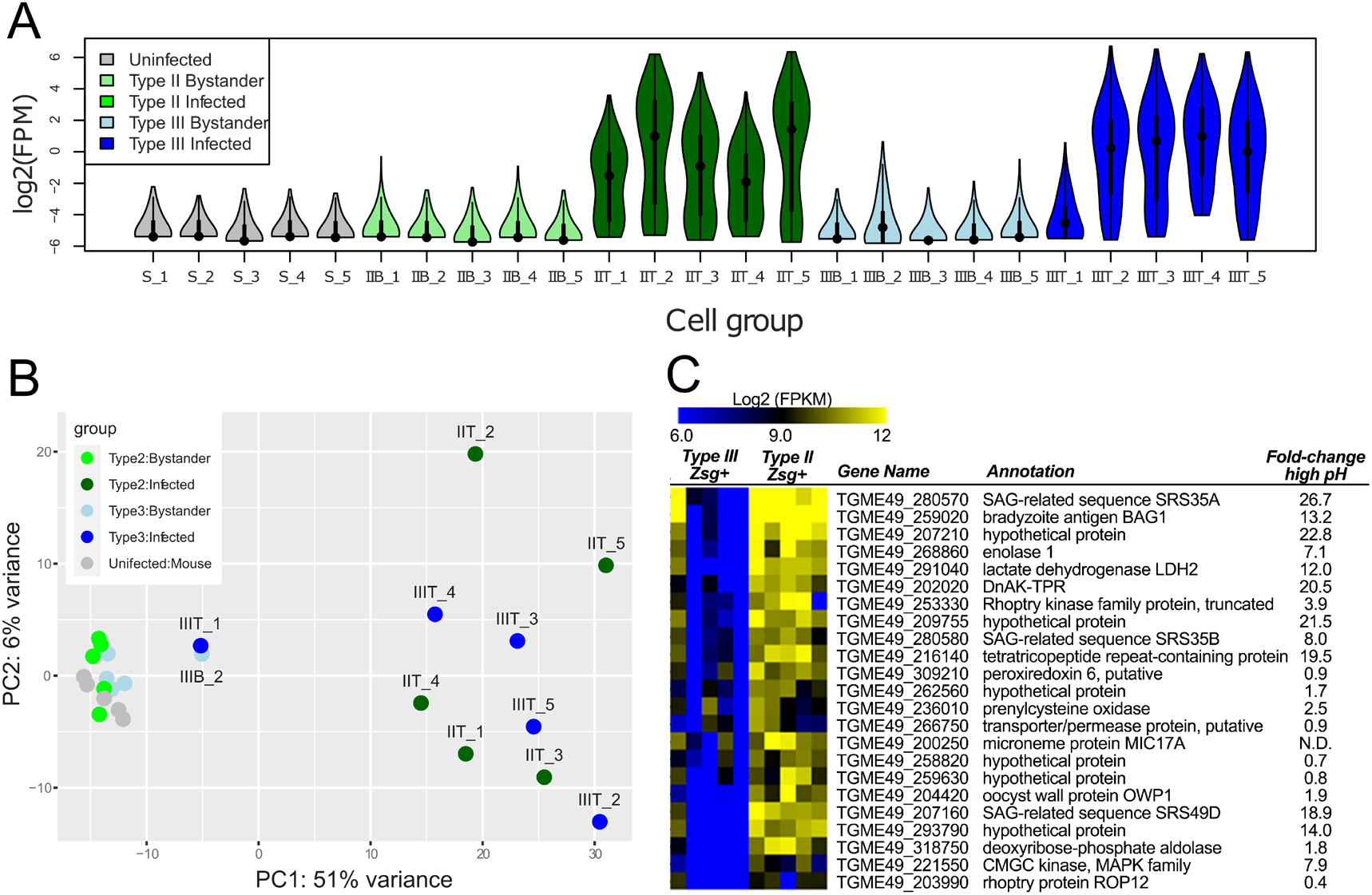
*T. gondii* bradyzoite transcripts have a higher abundance in type II TINs trancriptomes. **A)** Violin plot showing normalized (log2 FPM) read counts for *T. gondii* genes across samples. These graphs show data for 527 genes selected based on having fewer than 20 mapping reads across the 5 saline injected mouse control samples and at least 1 read in a minimum of 15 of the 25 samples. **B)** Principal components analysis of the data shown in **A**, illustrating general clustering between samples from Bystander cells and saline inoculated controls, and separation between TINs and all other samples. **C)** Log2 FPM values for a cluster of genes with distinct transcript abundance between type II and type III TINs. This cluster includes multiple canonical bradyzoite markers including BAG1, DNak-TPR and LDH2. In addition, most genes in this cluster are known to increase in abundance after high pH exposure in multiple *T. gondii* strain types, including the type III strain VEG^52^.

To look at only the infected cells in isolation from the Bystander and cells from uninfected mice (Saline controls), we first selected all genes with at least 1 read in at least 6 of the 25 samples (leaving 1,927 genes) and used DESeq2 to normalize and transform the data. We performed cluster analysis and found that most genes were similarly expressed between strains (**Fig. S3A**). While few genes were found to be significantly different in abundance between strains (likely due to the high variance in transcript counts across samples), we did identify a cluster of genes that were consistently of higher transcript abundance in type II TINs compared to type III TINs (**Fig. 5C**). Genes within this cluster included multiple canonical markers for the bradyzoite life stage, including BAG1, LDH2 and DnAK-TPR (**Fig. S3 and 5C**), suggesting a possible difference in bradyzoite status between type II and type III strains at the D21 timepoint when these samples were collected. To further explore these data, we compared the cluster of *T. gondii* genes with higher abundance in type II TINs versus type III TINs transcriptomes with published transcript abundance from a different type III strain (VEG) after bradyzoite conversion via high pH (**Fig. 5C**; data taken from Sokol et al.^52^). In this comparison, we found that many of the genes in the higher-in-type II TINs cluster were also induced in VEG bradyzoites compared to tachyzoites. When we performed Gene Set Enrichment Analysis (GSEA)^53^ using the previously published data^52^ to create “bradyzoite” and “tachyzoite” gene sets, we found significant enrichment of the bradyzoite gene set in type II TINs transcriptome and the tachyzoite gene set in type III TINs transcriptome (**Fig. S3B**). Overall these data are consistent with a difference in bradyzoite transcript abundance, and possibly developmental status, between type II and III strains at this time point.

## Discussion

Here we sought to gain the first insights into how *T. gondii* parasites manipulate neurons *in vivo* by leveraging a mouse model in which parasites trigger host cell green fluorescent protein (GFP) expression^34,54^ in combination with laser capture microdissection and RNAseq. We chose to use LCM to isolate both *T. gondii*-injected neurons (TINs) and nearby un-injected neurons (Bystanders) to control for paracrine cytokine effects that do not occur in uninfected mice. Using this technique, we generated high quality host and parasite transcripts that showed robust group-specific transcriptional profiles (**Fig. 3A**, **S2**) despite the relatively small sample size (200 cells/group). Unexpectedly, the separation between the groups, including between TINs and Bystanders, was driven primarily by differences in immune cell genes rather than differences in neuron signaling pathways (**Fig. 4A**). Further analysis suggested that immune cell clustering around TINs, and, to a lesser degree, Bystander neurons, explained much of the group-specific differences (**Fig. 4B**). In addition, a complex parasite transcriptional profile was detected in TINs and this profile was notably absent from Bystander transcriptomes. Collectively, these data suggest that at least a subpopulation of the TINs that we collected were infected with *T. gondii* parasites (in addition to being injected) and that immune cells, especially CD8 T cells, hone-in on TINs.

Why is our identification of immune cell transcripts in the *T. gondii*-infected brain novel? All prior genome-wide expression studies of the *T.gondii*-infected CNS have also identified high levels of immune cell or immune response transcripts compared to uninfected brain^48,49,51^. But these studies have all been done using whole brain for RNA isolation, where one would expect to obtain transcripts from all cells within the brain (parenchymal CNS cells and infiltrating immune cells). Conversely, we tried to avoid non-neuronal cells by using laser capture microdissection to isolate 10 μm diameter areas centered on neuron somas, which are ~10 μm in diameter. In addition, instead of simply comparing our TINs transcriptomes to transcriptomes derived from uninfected mice, we isolated bystander neurons which are 8-12 cell bodies away from an isolated TIN. The enrichment for immune cell transcripts in the TINs transcriptomes compared to transcriptomes from uninfected mice and Bystander neurons indicates that these immune cells, especially CD8 T cells, are in close proximity to TINs, suggesting the T cells may “recognize” the TINs. The potential recognition of infected or injected neurons by T cells supports a radical shift in our understanding of neuron capabilities for generating cellular immune responses. Only in the last several decades have neurons been shown to express MHC I at baseline and, *in vitro*, have the capability to stimulate CD8^+^ T cells^55–58^. An increase in T cell transcripts from bits of immune cells being captured in close proximity to LCM-captured TINs compared to Bystanders, in conjunction with the presence of *T. gondii* transcripts in TINs, further strengthens the possibility an interaction between neurons, *T. gondii*, and T cells. One caveat, though, is that because our individually isolated cells were pooled (i.e. all 200 TINs from a single mouse were collected together after which RNA was isolated), we cannot distinguish whether the immune cell signatures arose equally from each TIN or if a subset of TINs had much higher numbers of aggregated immune cells (**Fig. 6**). While future work will focus on distinguishing between these possibilities, either model adds to the growing literature that, contrary to dogma, infected neurons have robust immune responses to microbes and cytokines and can present antigens to T cells *in vivo*^7,59,50^.

**Figure 6.**
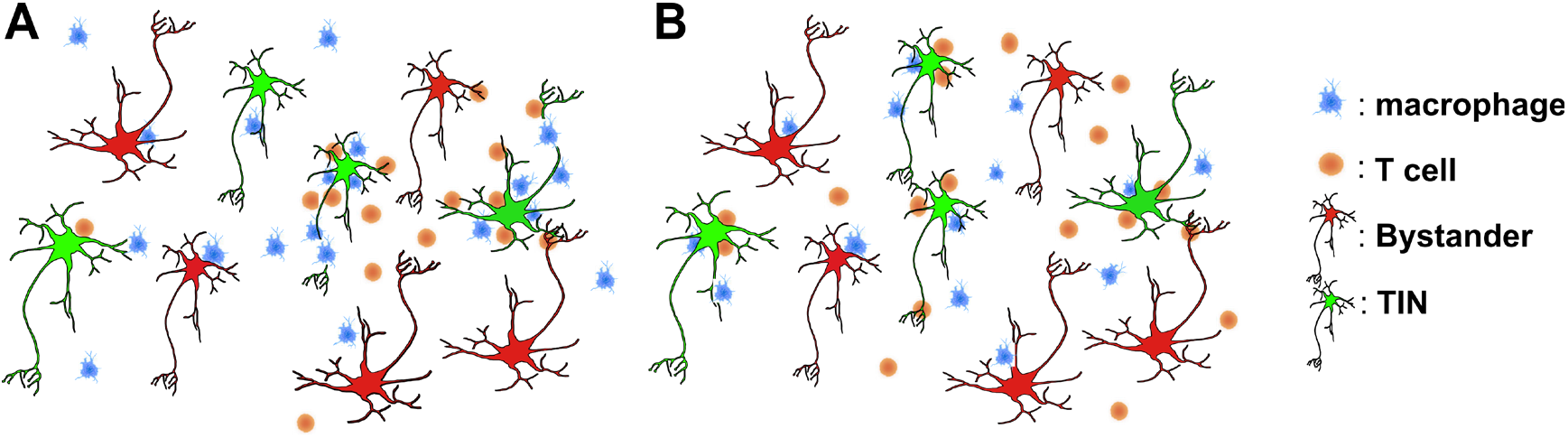
Proposed models of immune cell clustering around TINs. **A)** Model of macrophages and T cells clustering around select TINs. **B)** Model of uniform clustering of macrophages and T cells around TINs

One complication of the high abundance of immune transcripts is that we were unable to accomplish our original goal of defining *T.gondii* strain-specific (II-Cre and III-Cre) manipulations of neurons. When normalized to saline, the top genes in II-Cre TINs and III-Cre TINs are almost identical and largely composed of immune response transcripts (**Fig. S4**). Even when we only compare III-TINs to II-TINs, which leads to strain-specific separation of samples on PCA analysis, we are unable to distinguish what transcripts are inherent to neurons and what transcripts are from clustering immune cells (**Fig. S4**). Thus, future work to define *T. gondii* strain-specific differences in neuron signaling will require a technique in which neurons can be cleanly separated from the infiltrating immune cells, such as brain digestion followed by cell sorting. Such a technique was used in the only other study that has sought to identify how microbe-manipulated neurons are affected *in vivo*^60^. This study used an attenuated rabies virus-Cre system in combination with flow cytometry to isolate previously infected neurons at 6 months post infection^60^. While that study was able to successfully separate out RBV-affected neuron transcripts, as the uninfected, bystander neurons came from throughout the mouse brain, it could not control for paracrine changes versus direct RBV-induced changes. Nor could the study be used to identify RBV transcripts as the time point assessed was well beyond when the virus was cleared.

Conversely, we were able to identify parasite transcripts in the TINs transcriptomes (**Fig. 5A**). Akin to the immune cell transcripts, several models could account for the parasite transcripts being in the TINs transcriptomes, and future work will focus on determining if these transcripts arise from TINs (implying the TINs were infected at some time prior to harvest) versus other possible sources. Regardless of the origins of the transcript, the strain-specific differences in developmental transcripts— a higher abundance of bradyzoite genes in type II TINs versus a higher abundance of tachyzoite transcripts in type III TINs (**Fig. 5C**, **S3**)— is intriguing. These data suggest that type II and type III strains may have different rates of conversion from tachyzoites to bradyzoites *in vivo*, with the type III strain trailing behind the type II strain at the D21 timepoint, despite the fact that type III strains are very capable of forming cysts *in vivo*^23,32,61^. We recently described how the host immune response at the same time point differs between these strains^32^, leading to a proverbial chicken-or-the-egg question. Do strain-specific differences in host immune signals dictate the rate of parasite conversion upon arrival to the brain? Or do differences in the intrinsic rates of parasite stage conversion in the CNS leading to strain-specific immune responses? Or both? Ultimately, these data suggest that how T. gondii successfully establishes a persistent CNS infection may vary by strain type.

In summary, the data presented here suggest that neurons injected with *T. gondii* proteins are surrounded by infiltrating immune cells and that, *in vivo*, *T. gondii* strains may differ in the rate in which they convert from the lytic tachyzoite form to the persistent bradyzoite form. Collectively, these data suggest we have much to learn about neuron-parasite-immune cell interactions and how these interactions vary by *T. gondii* strain.

## Methods

### Ethics statement

All mouse studies and breeding were carried out in strict accordance with the Public Health Service Policy on Human Care and Use of Laboratory Animals. The protocol was approved by the University of Arizona Institutional Animal Care and Use Committee (#A-3248-01, protocol #12–391).

### Mice

All mice used in this study are Cre reporter mice that only express GFP in their cells after Cre-mediated recombination^33^. Mice were purchased from Jackson Laboratories (stock # 007906) and bred in the University of Arizona, BIO5 Animal Facility. Mice were inoculated intraperitoneally (i.p.) with freshly syringe-released parasites, diluted to the appropriate inoculums in 200 μl volume in USP grade PBS. The inoculating number of parasites was 10,000 (II-Cre) or 10,000 (III-Cre).

### Parasites

All strains were maintained through serial passage in human foreskin fibroblasts (gift John Boothroyd, Stanford University, Stanford, CA) using DMEM, supplemented with 10% fetal bovine serum, 2mM glutagro, and 100 I.U./ml penicillin/ 100 μg/ml streptomycin. Type II, Prugniaud, referred to as II-Cre, and type III, CEP, referred to as III-Cre, strains express Cre and mCherry^34^.

### LCM Brain Tissue preparation

Cre reporter mice were infected with an inoculum of 10,000 parasites for 21 days. Upon harvest, brains were removed and cut into 2 hemispheres. Brains were washed in sterile PBS containing ProtectRNA (Sigma Aldrich R7397-30ML), moved to a dish containing sterile 4% PFA for 5-10 minutes, and then transferred to another dish containing sterile PBS containing ProtectRNA. Hemispheres were then flash frozen in OCT and isopentane and sectioned to 8 um on Cryostat.

### LCM Brain Tissue Staining

Mounted sections were quickly thawed and dipped in nuclease free water and ProtectRNA (RNase Inhibitor; Sigma-Aldrich R7397-30ML) several times to remove O.C.T. Excess water was removed from the slides and 100-200 μl antibody solution was added to the sections (Anti-NeuN Antibody, clone A60, Alexa Fluor 555 Conjugate – MAB377A5 (1:200)) and placed on cold plate on ice, covered, for 10 minutes. Excess antibody was wicked away and sections were washed two times with sterile PBS and ProtectRNA for 10 seconds. Samples were dehydrated by consecutively submerging slides in 70% EtOH for 60 seconds, 95 % EtOH for 60 seconds and 100% EtOH for 60 seconds. 100% EtOH step was repeated two times, followed by two 60 second washes in xylenes^62^. Samples were air dried for 5 minutes and LCM was performed immediately.

### LCM

Laser capture microdissection was performed on Arcturus XT Laser Capture Microdissection system with Nikon Eclipse Ti-E Microscope Base. The Arcturus XT Epifluorescent Illumination Package utilizes custom filter cubes: green: excitation 503 nm-548 nm, emission >565 nm, red:excitation 570 nm - 630 nm, emission > 655nm, and triple Dichroic cube (DAPI/FITC/TRITC) (part # 6530-0056) excitation 385-400/ 475-493/ 545-565 nm, emission 450-465/ 503-533/ 582-622 nm.

### RNA isolation, preparation of cDNA libraries and sequencing

RNA Samples were assessed for quality with an Advanced Analytics Fragment Analyzer *(High Sensitivity RNA Analysis Kit # DNF-491)* and quantity with a Qubit RNA quantification kit *(Qubit® RNA HS Assay Kit # Q32855)*. Samples were used for library builds with the ClonTech SMART-Seq V4 Ultra Low Input RNA Kit from Takara *(Catalog # 634890)*. Upon library build completion, samples had quality and average fragment size assessed with the Advanced Analytics Fragment Analyzer *(High Sensitivity NGS Analysis Kit # DNF-486)*. Quantity was assessed with an Illumina Universal Adaptor-specific qPCR kit from Kapa Biosystems *(Kapa Library Quantification kit for Illumina NGS # KK4824)*. After final library QC was completed, samples were equimolar-pooled and clustered for sequencing on the NextSeq500 machine. The paired-end sequencing run was performed using Illumina HighSeq2500 run chemistry *(HiSeq2500 High Output 2×100PE 8-lane run, 200 total cycles, FC-401-3001)*.

### Data processing and differential gene expression analysis, host transcripts

An average of 57.7 million raw reads per library were obtained for 25 distinct barcoded Illumina RNAseq libraries (5 control, 5 TINs II-Cre, 5 Bystander II-Cre, 5 TINs III-Cre, and 5 Bystander III-Cre). To prepare the raw data for differential gene expression analysis, the reads from the Illumina FASTQ files from each Illumina library were pseudoaligned to the *Mus musculus* reference transcriptome (ENSEMBL GRCm38.81) and a determination of transcript abundance was performed with Kallisto, which utilizes a combination of k-mer hashing and a transcriptome de Bruijn graph for accurate pseudoalignment^36^. The transcript-level abundance measures that were determined by Kallisto were converted to length-scaled gene-level abundances with tximport and reported in transcripts per million reads (TPM) units^63^. The length-scaled gene-level abundances were then analyzed for differential gene expression between the 5 analysis groups utilizing DESeq2^35^. Tximport and DESeq2 analysis were performed on the CyVerse infrastructure^64^.

The analysis was performed using the R software^65^, Bioconductor^66^ packages including DESeq2^67,35^ and the SARTools package developed at PF2 - Institut Pasteur. Normalization and differential analysis are carried out according to the DESeq2 model and package. Briefly, the raw data was reported with mapped reads per group, the proportion of null reads not included in further analysis, and the distribution of reads across groups which were very similar. To assess the similarity between samples within replicates and across conditions, the Simple Error Ratio Estimate (SERE) statistic was used to measure whether the variability between samples is random Poisson variability or higher^68^. DESeq2 then transforms the data to account for differential variance across range of the means to render data that approaches homoscedasticity, using the Variance Stabilizing Transformation (VST)^35,67^ method as shown in cluster dendrogram and PC plots. DESeq2 then normalized the data by computing a scalar factor for each sample, assuming that most of the genes will not be differentially expressed, with the default setting locfunc = “median”. The differential analysis performed by DESeq2 fits one linear model per feature with log_2_(FC) that calculated p and q values. Outliers were calculated by Cook’s distance^69^ and were not assigned a p-value. The dispersions estimate was set to the default setting of Generalized Linear Model (GLM). Then, DESeq2 imposed a Cox Reid-adjusted profile likelihood maximization^70,71^ and used the maximum a posteriori (MAP) of the dispersion^72^. DESeq2 finally plots the raw p-values for differential expression and performs independent filtering to increase detection power^35^. For the final results, including an adjusted p-value calculation, a Benjamini-Hochberg (BH) p-value adjustment was performed^73,74^ and the level of controlled false positive rate was set to 0.05. Raw sequencing data will be available on ToxoDB (https://toxodb.org/toxo/), which now a part of VEuPathDB.org.

### Pathway Analysis

Differentially expressed genes were uploaded into Ingenuity Pathway Analysis (Qiagen), with one set normalized to saline as well as TIN groups normalized to their respective Bystanders (type II TINs normalized to type II Bystanders, etc). A “Core Analysis” was run on each group before a “Comparison Analysis” was run between groups. “Canonical Pathways” and “Diseases and Functions” analysis was performed for inflammatory and immune signals with the following filter settings: Canonical pathways –> Filter -> Cellular Immune Response, Cytokine Signaling (T Helper Cell Differentiation, Th1 Pathway, Th1 and Th2 Activation Pathways, Th2 Pathway), Humoral Immune Response, and Pathogen-Influenced Signaling. Diseases and Functions -> Filter -> Diseases and Disorders (Antimicrobial Response), Molecular and Cellular Functions (Cell-to-Cell Signaling, Cellular Function and Maintenance, and Cellular Movement), and Molecular and Cellular Functions, Physiological System Development and Function.

### CIBERSORT analysis

Normalized gene expression values for uninfected, bystander, and TIN samples and the LM22 signature matrix were used as input to CIBERSORT^40^. The deconvolution was performed on Stanford University’s online CIBERSORT webpage tool. Both absolute and relative modes were run, and quantile normalization was disabled. 1000 permutations were run for statistical testing. Saline samples had p-values = 1, suggesting that small levels (< 4%) of immune cell abundance in the saline samples were not significant. For this reason, for samples with p-value <0.05, only immune cell populations above the saline based threshold of 4% are shown.

### Data processing and differential gene expression analysis, parasite transcripts

Fastq files were first mapped against the mouse transcriptome (GRCh38 v21; parameters −k 5, --very-sensitive-local) and the non-mapping reads then mapped to the *T. gondii* ME49 genome (v44; parameters −k 5 −very-sensitive-local). Raw transcript counts were determined using featureCounts from the Subread package integrating the gff file from ToxoDB and the sorted bam file for each sample. Genes were removed from the analysis if they had ≥ 20 total mapping reads across the 5 LCM samples obtained from saline-injected mice. Genes were then normalized in two different ways. To determine the relative quantity of parasite-specific reads across all samples, data were transformed using the formula log_2_(1×10^6*^(read counts/total host mapping reads)) where total mapping reads were obtained from **Table S1** to generate the log_2_(FPM) value and then filtered based on read counts across samples to include only those genes with at least 1 read in at least 15 of the 25 samples (resulting in 527 genes total). For analysis of the TINs separately from Bystanders and cells from mock-treated control mice, data were included only if they had at least 1 read in 6 of the 25 samples (resulting in 1927 genes total), and raw counts were uploaded into the DESeq2 package in R and normalize using the “rlog” function. Differential expression between strains was determined using default settings (“results”). These two methods of normalization are why the log_2_(FPM) values are of different scales.

## Acknowledgments

The authors would like to thank the whole Koshy laboratory for fruitful discussions. The author(s) disclosed receipt of the following financial support for the research, authorship, and/or publication of this article: Funding by the National Institutes of Health [R01NS095994 (A. A. K); R01HL147187 (C.E.R); R01AI116855 (J.P.B.); T32-AG058503 (E. F. M).; T32-AG061897 (H. J. J.); T32-HL007249 (A.C.C)]; the Achievement Rewards for College Scientists (ARCS) Foundation in Phoenix (A.C.C., H. J. J.); the March of Dimes [#5-FY15-45 (AAK)]; the BIO5 Institute, University of Arizona (A. A. K.); and the National Science Foundation [DBI-0735191, DBI-1265383,and DBI-1743442 (www.cyverse.org)]. The funders had no role in study design; data collection, analysis, or interpretation; or the decision to submit the work for publication.

