## Supplemental Figures 1-4 and Tables 1 and 4 for "Transcriptional profiling reveals T cells cluster around neurons injected with *Toxoplasma gondii* proteins"

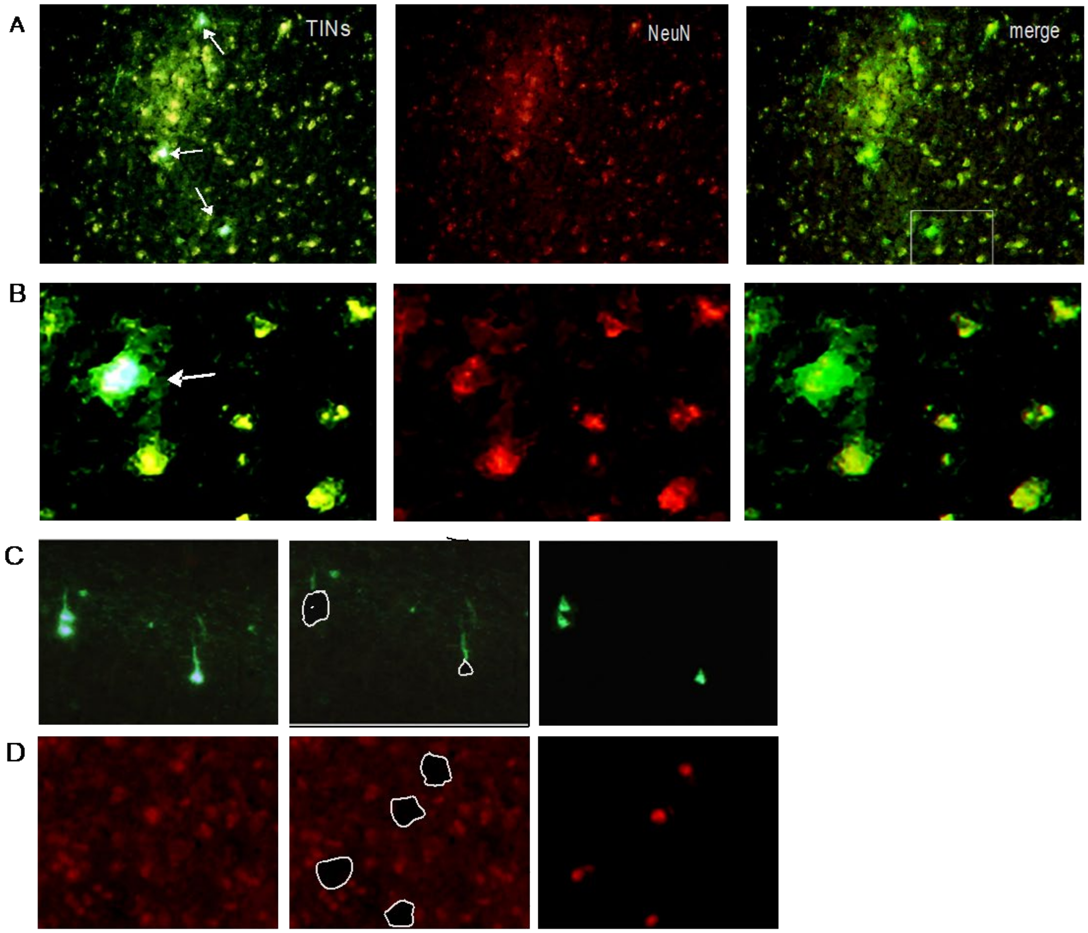


**Figure S1. Isolation of soma by laser capture microdissection (LCM). A)** TINs (white arrows) in 8 micron thick brain section imaged on Arcturus XT Laser Capture Microdissection system with Nikon Eclipse Ti-E Microscope base. Only white arrows are TINs, residual green stain is due to microscope filters. Microscope uses a mercury lamp, and filter cubes that excite (green) 503 nm -548 nm and (red) 570 nm – 630 nm. We used a NeuN conjugated antibody to 555 fluorophore. Therefore, when we look in the green channel to visualize TINs, a level of $NeuN^+$ 555 antibody is also excited and visualized. In the green channel, the difference between $GFP^+$ $NeuN^+$ cells and cells that were $NeuN^+$ 555 was apparent. Panel 2, $NeuN^+$ staining shows neuronal cell body as viewed under microscope. Panel 3, merged image shows TINs are neurons. **B)** Enlargement of A. **C)** Example of soma isolated by LCM. Panel 1 shows original image. Panel 2 shows brain section after isolation of soma, area removed is outlined in white. Panel 3 shows soma on cap after isolation. Bleedthrough of $NeuN^+$ staining is not present in C because these are test sections, not stained with NeuN antibody. **D)** $NeuN^+$ staining of brain section. Panel 1 shows original image. Panel 2 shows brain section after retrieval of soma. Panel 3 shows isolated soma on cap.

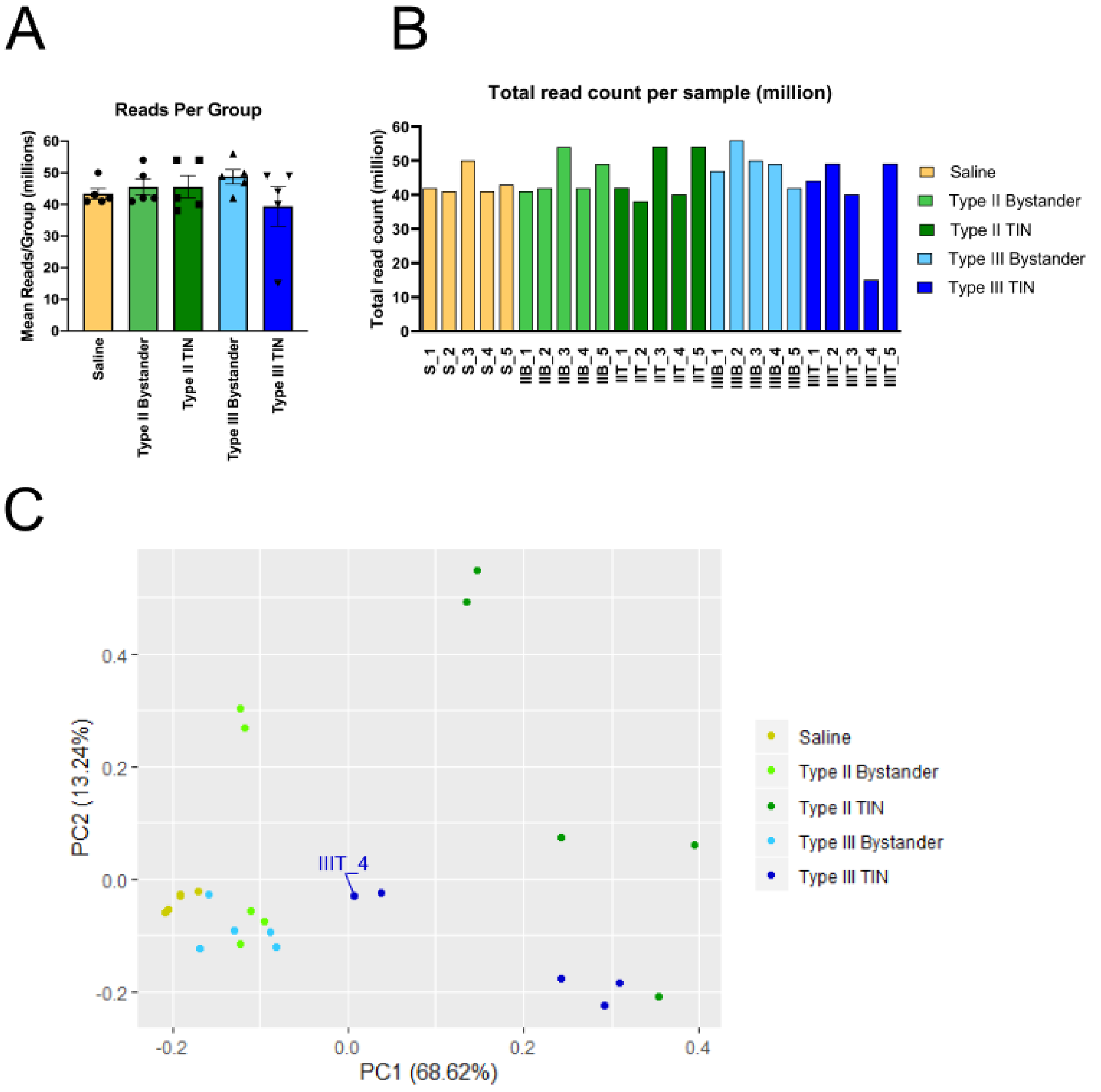


**Figure S2. Reads are almost entirely consistent within and across samples**. **A)** Mean number or reads per group. **B)** Total number of mapped reads per sample. Colors refer to the biological condition of the sample. Reads that map on multiple locations on the transcriptome are counted more than once, as far as they are mapped on less than 50 different loci. Total read counts are similar within and across conditions with IIIT_4 as the one exception, though IIIT_4 still mapped appropriately (see **S4**). **C)** Rounded counts from Kallisto were analyzed in R Studio. PC1 accounts for 68.62% of the variation, while PC2 accounts for 13.24% of the variation. The TINs are separated from the Bystanders and saline by both principle components, revealing the most variation within the dataset. IIIT_4 groups with another type III TIN in transcript variability.

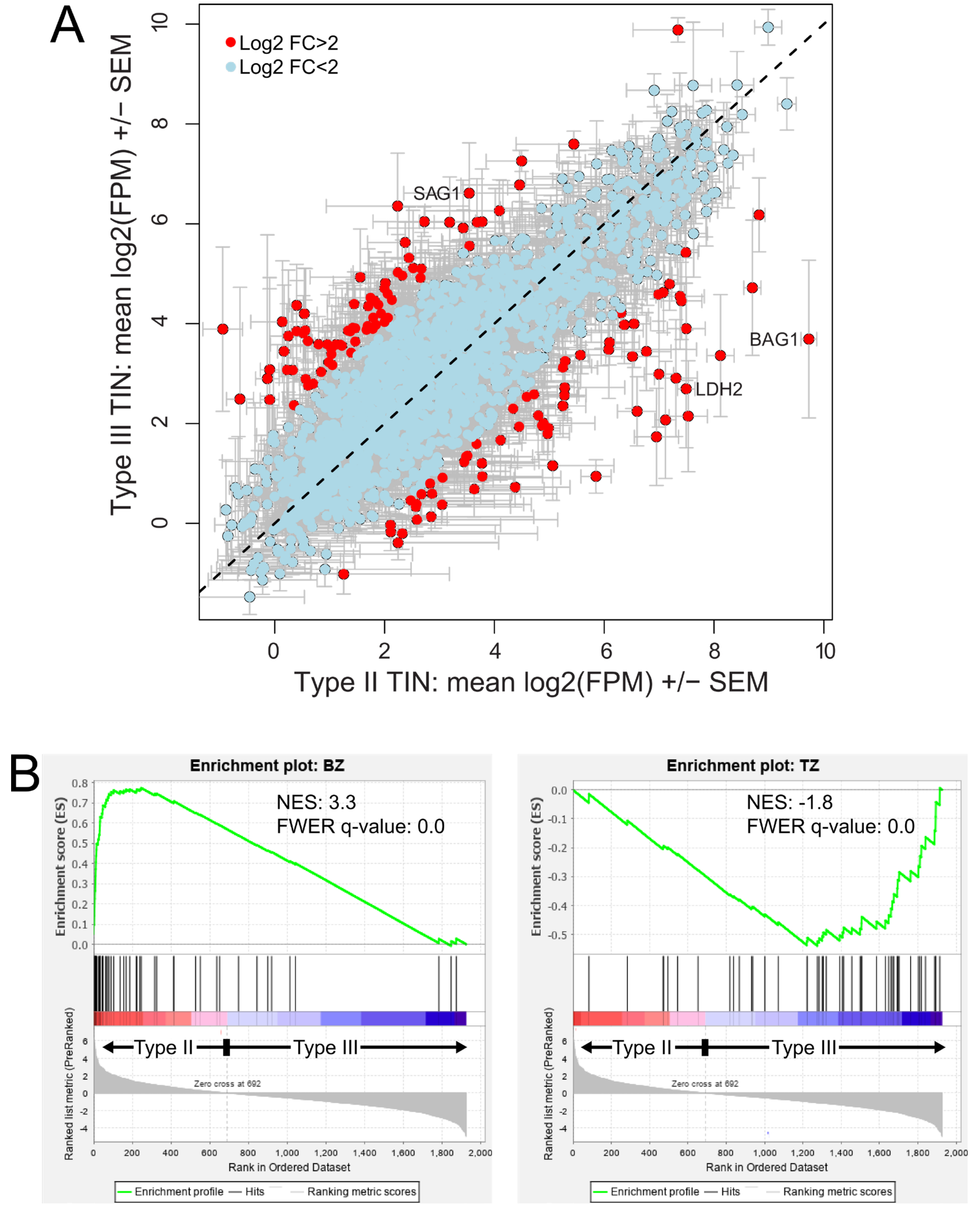


**Figure S3. Type II and type III *T. gondii* strains differ in bradyzoite and tachyzoite gene expression. A)** Plot comparing $log_2$ (FPM) values for 1,923 *T. gondii* transcripts identified for type II and type III TINS. Data were selected if reads could be found in at least 6 of the 25 samples. Error bars represent SEM (N=5 for each strain type), and datapoints are colored red if they had at least a $log_2$ difference of 2 between type II or type III TINS. **B)** Gene Set Enrichment Analysis of 1,923 transcripts ranked by transcript abundance in type II versus type III TINS compared to a curated bradyzoite (left) and tachyzoite (right) gene set. Type II parasites have significant enrichment for the BZ gene set while type III parasites have significant enrichment for the TZ gene set.

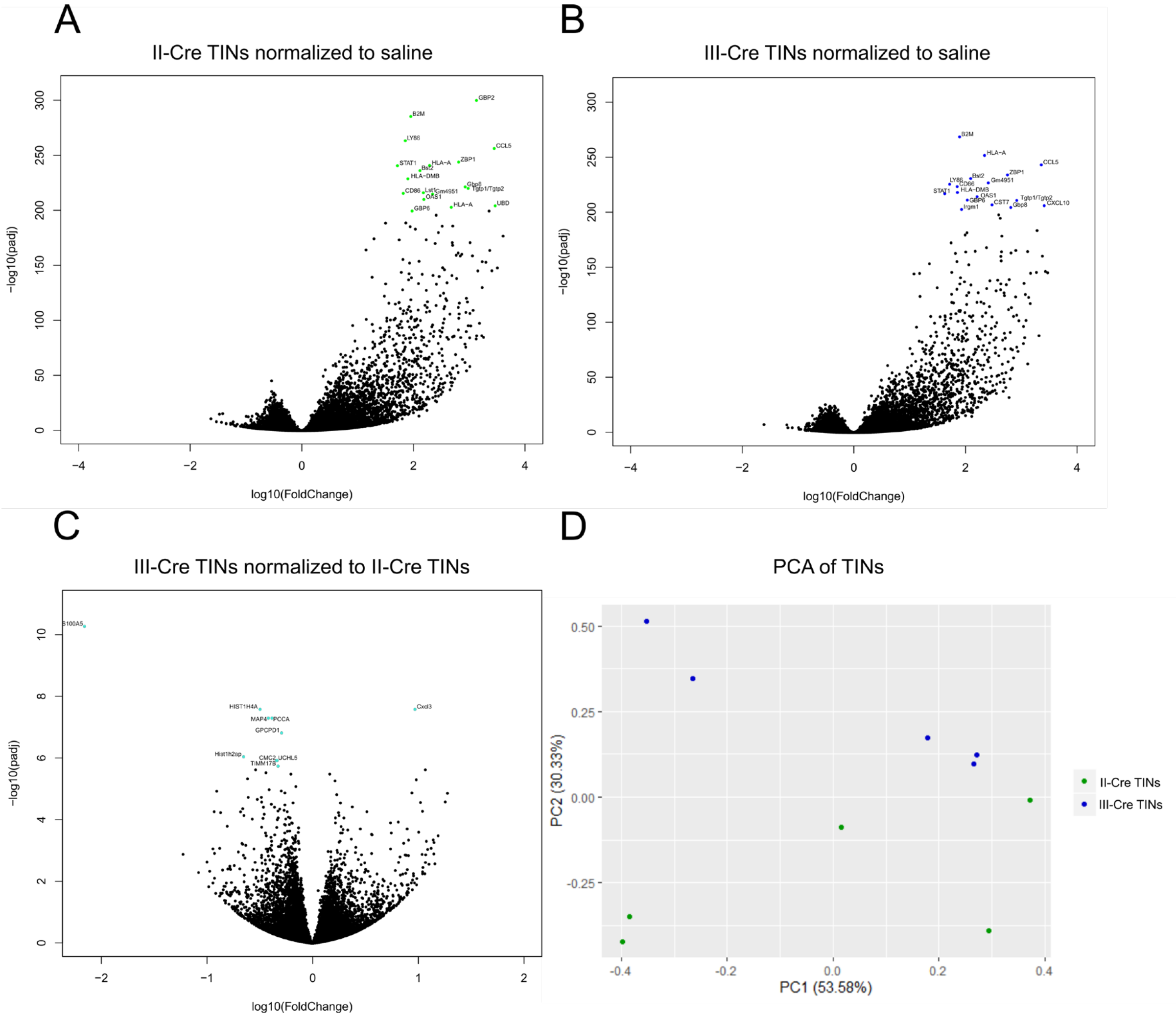


**Figure S4. The most upregulated transcripts in Type II and Type III TINs are immune related.** DEGs from DESeq2 for II-Cre TINs and III-Cre TINs normalized to saline were analyzed in R Studio. The top 12 genes are highlighted in **A)** green for II-Cre TINs and **B)** blue for III-Cre TINs. Many of the same genes are expressed in both groups and are immunity related. DEGs from DESeq2, with III-Cre TINs normalized to II-Cre TINs, reveal differences in transcripts are possibly due to immune cells. **C)** Volcano plot of type III TINs normalized to type II TINs. As an example, S100A5, downregulated in III-Cre TINs compared to II-Cre TINs in the upper left corner, may be due to infiltrating immune cells and not neuronal response. **D)** PCA plot of TINs. Principle Component 2 (PC2) separates TINs by strain. Type II TINs cluster to top of graph while type III TINs are only in bottom half of graph.

| Sample Number | Number of Reads | Number of Mapped Reads | Sample Type | Renamed Sample # |
|---|---|---|---|---|
| 1 | 61,236,295 | 42,091,657 | Saline | S_1 |
| 2 | 53,230,580 | 41,559,705 | Saline | S_2 |
| 3 | 67,508,516 | 50,760,718 | Saline | S_3 |
| 4 | 53,775,849 | 41,761,779 | Saline | S_4 |
| 5 | 55,744,725 | 43,668,262 | Saline | S_5 |
| 6 | 56,715,483 | 42,188,554 | Type II Bystander | IIB_1 |
| 7 | 55,864,901 | 43,391,341 | Type II Bystander | IIB_2 |
| 8 | 68,437,547 | 53,333,496 | Type II Bystander | IIB_3 |
| 9 | 58,790,086 | 43,610,986 | Type II Bystander | IIB_4 |
| 10 | 63,495,652 | 49,239,523 | Type II Bystander | IIB_5 |
| 6 | 53,948,729 | 43,085,864 | Type II TINS | IIT_1 |
| 7 | 49,829,178 | 39,386,921 | Type II TINS | IIT_2 |
| 8 | 61,443,459 | 49,080,858 | Type II TINS | IIT_3 |
| 9 | 52,974,279 | 41,922,336 | Type II TINS | IIT_4 |
| 10 | 68,049,310 | 53,586,903 | Type II TINS | IIT_5 |
| 11 | 56,300,608 | 46,412,206 | Type III Bystander | IIIB_1 |
| 12 | 69,272,836 | 56,144,674 | Type III Bystander | IIIB_2 |
| 13 | 59,712,525 | 49,499,213 | Type III Bystander | IIIB_3 |
| 14 | 58,555,830 | 48,373,523 | Type III Bystander | IIIB_4 |
| 15 | 53,307,102 | 43,360,355 | Type III Bystander | IIIB_5 |
| 11 | 58,518,851 | 45,803,170 | Type III TINS | IIIT_1 |
| 12 | 63,441,555 | 48,388,826 | Type III TINS | IIIT_2 |
| 13 | 55,807,398 | 42,253,369 | Type III TINS | IIIT_3 |
| 14 | 22,121,262 | 16,439,167 | Type III TINS | IIIT_4 |
| 15 | 63,415,869 | 48,742,844 | Type III TINS | IIIT_5 |

**Table S1. The number of reads per FASTQ RNAseq file have narrow variation within and across samples.** Sample number corresponds to the mouse from which samples were taken from. Bystanders and TINs were captured from the same mice. The number of reads per FASTQ RNAseq file have narrow variation within and across samples. The renamed sample numbers are the names that will often be used throughout figures.

| Input Sample | B cells naive | B cells memory | Plasma cells | T cells CD8 | T cells CD4 naive | T cells CD4 memory resting | T cells CD4 memory activated | T cells follicular helper | T cells regulatory (Tregs) | T cells gamma delta | NK cells resting | NK cells activated | Monocytes | Macrophages M0 | Macrophages M1 | Macrophages M2 | Dendritic cells resting | Dendritic cells activated | Mast cells resting | Mast cells activated | Eosinophils | Neutrophils | P-value | Pearson Correlation | Absolute score |
|---|---|---|---|---|---|---|---|---|---|---|---|---|---|---|---|---|---|---|---|---|---|---|---|---|---|
| S_1 | 1.9% | 0.0% | 1.4% | 0.5% | 0.7% | 0.0% | 0.0% | 0.3% | 0.0% | 0.9% | 0.0% | 0.1% | 0.0% | 0.0% | 0.0% | 0.0% | 1.9% | 0.0% | 0.0% | 0.0% | 0.0% | 0.6% | 1.000 | -0.095 | 0.082 |
| S_2 | 1.7% | 0.0% | 2.1% | 0.0% | 0.0% | 0.9% | 0.0% | 0.3% | 0.0% | 0.0% | 0.0% | 1.0% | 0.0% | 0.1% | 0.0% | 0.8% | 0.0% | 0.0% | 0.7% | 0.0% | 0.0% | 0.3% | 1.000 | -0.107 | 0.079 |
| S_3 | 3.0% | 0.0% | 1.6% | 0.0% | 1.1% | 0.0% | 0.0% | 0.5% | 0.0% | 0.6% | 0.0% | 0.4% | 0.0% | 0.0% | 0.0% | 0.8% | 1.3% | 0.0% | 0.2% | 0.0% | 0.0% | 0.0% | 1.000 | -0.092 | 0.096 |
| S_4 | 1.5% | 0.0% | 0.6% | 0.0% | 0.6% | 0.0% | 0.0% | 0.3% | 0.0% | 0.0% | 0.0% | 0.2% | 0.0% | 0.0% | 0.0% | 0.4% | 1.1% | 0.0% | 0.0% | 0.0% | 0.4% | 0.0% | 1.000 | -0.115 | 0.049 |
| S_5 | 1.7% | 0.0% | 1.0% | 0.0% | 0.2% | 0.0% | 0.0% | 0.4% | 0.0% | 0.0% | 0.0% | 0.2% | 0.0% | 0.0% | 0.0% | 0.4% | 1.4% | 0.0% | 0.0% | 0.0% | 0.0% | 0.0% | 1.000 | -0.084 | 0.053 |
| Average_S | 2.1% | 0.0% | 1.5% | 0.0% | 1.1% | 0.0% | 0.0% | 0.8% | 0.0% | 0.0% | 0.0% | 0.2% | 0.0% | 0.0% | 0.0% | 0.9% | 0.7% | 0.0% | 0.0% | 0.0% | 0.3% | 0.0% | 1.000 | -0.085 | 0.076 |
| IIB_1 | 1.3% | 0.0% | 0.0% | 11.8% | 0.0% | 0.0% | 0.0% | 0.6% | 0.0% | 0.0% | 0.0% | 1.2% | 3.0% | 0.0% | 2.2% | 1.7% | 0.7% | 0.0% | 1.5% | 0.0% | 0.0% | 0.0% | 0.020 | 0.253 | 0.238 |
| IIB_2 | 2.7% | 0.0% | 0.0% | 12.5% | 0.0% | 0.0% | 1.3% | 1.3% | 0.0% | 0.0% | 0.0% | 1.1% | 4.1% | 0.0% | 3.9% | 1.7% | 1.1% | 0.1% | 2.2% | 0.0% | 0.0% | 0.0% | 0.000 | 0.363 | 0.319 |
| IIB_3 | 1.9% | 0.0% | 0.0% | 17.5% | 0.0% | 0.0% | 0.0% | 0.0% | 0.0% | 2.1% | 0.0% | 0.0% | 7.3% | 0.0% | 4.3% | 3.8% | 2.6% | 0.0% | 0.7% | 0.0% | 0.0% | 0.0% | 0.000 | 0.338 | 0.402 |
| IIB_4 | 1.4% | 0.0% | 0.0% | 10.3% | 0.0% | 0.0% | 0.0% | 0.1% | 0.1% | 1.9% | 0.0% | 0.7% | 2.3% | 0.0% | 1.9% | 2.3% | 0.0% | 0.0% | 2.1% | 0.0% | 0.2% | 0.0% | 0.020 | 0.258 | 0.234 |
| IIB_5 | 2.0% | 0.0% | 0.0% | 19.5% | 0.0% | 0.0% | 0.0% | 0.6% | 0.0% | 0.0% | 0.0% | 2.3% | 5.5% | 0.0% | 4.3% | 2.7% | 2.5% | 0.0% | 1.2% | 0.0% | 0.0% | 0.0% | 0.000 | 0.381 | 0.405 |
| Average_IIB | 3.3% | 0.0% | 0.0% | 14.3% | 0.0% | 0.0% | 0.7% | 0.2% | 0.9% | 0.0% | 0.0% | 1.7% | 5.8% | 0.0% | 4.1% | 1.7% | 1.3% | 0.0% | 2.0% | 0.0% | 0.0% | 0.0% | 0.000 | 0.332 | 0.359 |
| IIT_1 | 0.4% | 0.0% | 0.0% | 32.9% | 0.0% | 0.0% | 2.5% | 0.0% | 0.2% | 0.0% | 0.0% | 3.3% | 9.3% | 0.0% | 6.4% | 3.1% | 0.3% | 0.6% | 3.6% | 0.0% | 0.0% | 0.0% | 0.000 | 0.468 | 0.627 |
| IIT_2 | 0.0% | 0.0% | 0.2% | 22.2% | 0.0% | 0.0% | 3.3% | 0.0% | 1.2% | 2.2% | 0.0% | 0.3% | 7.7% | 0.0% | 4.4% | 1.5% | 1.7% | 0.0% | 1.4% | 0.0% | 0.0% | 0.0% | 0.000 | 0.429 | 0.461 |
| IIT_3 | 0.3% | 0.0% | 0.0% | 29.0% | 0.0% | 0.0% | 0.3% | 1.6% | 0.1% | 0.0% | 0.0% | 1.7% | 11.2% | 0.0% | 8.7% | 1.5% | 3.2% | 0.0% | 2.5% | 0.0% | 0.0% | 0.0% | 0.000 | 0.434 | 0.602 |
| IIT_4 | 0.0% | 0.0% | 0.0% | 32.1% | 0.0% | 0.0% | 1.3% | 1.3% | 0.0% | 0.0% | 0.0% | 4.3% | 10.1% | 0.0% | 5.6% | 3.3% | 0.4% | 1.4% | 3.0% | 0.0% | 0.0% | 0.0% | 0.000 | 0.450 | 0.628 |
| IIT_5 | 0.0% | 0.0% | 0.0% | 30.5% | 0.0% | 0.0% | 1.6% | 0.4% | 1.7% | 0.0% | 0.0% | 3.3% | 10.1% | 0.0% | 7.0% | 3.4% | 5.4% | 0.0% | 2.6% | 0.0% | 0.0% | 0.0% | 0.000 | 0.423 | 0.660 |
| Average_IIT | 0.4% | 0.0% | 0.0% | 32.2% | 0.0% | 0.0% | 2.0% | 1.0% | 0.0% | 0.0% | 0.0% | 1.7% | 9.7% | 0.0% | 6.8% | 2.3% | 3.4% | 0.0% | 2.4% | 0.0% | 0.0% | 0.0% | 0.000 | 0.439 | 0.620 |
| IIIB_1 | 0.8% | 0.0% | 0.0% | 9.8% | 0.0% | 0.0% | 2.8% | 0.0% | 0.0% | 1.7% | 0.0% | 0.3% | 6.0% | 0.2% | 1.1% | 1.9% | 0.0% | 0.0% | 1.1% | 0.0% | 0.0% | 0.0% | 0.030 | 0.211 | 0.256 |
| IIIB_2 | 1.5% | 0.0% | 0.0% | 28.1% | 0.0% | 0.0% | 1.1% | 0.0% | 0.1% | 0.0% | 0.0% | 0.5% | 10.2% | 0.0% | 1.0% | 5.7% | 0.9% | 0.0% | 1.1% | 0.0% | 0.0% | 0.0% | 0.020 | 0.290 | 0.502 |
| IIIB_3 | 2.5% | 0.0% | 0.0% | 9.7% | 0.0% | 0.0% | 0.0% | 0.8% | 0.0% | 0.0% | 0.0% | 1.5% | 4.1% | 0.0% | 0.7% | 3.1% | 0.0% | 0.0% | 2.3% | 0.0% | 0.0% | 0.0% | 0.130 | 0.117 | 0.247 |
| IIIB_4 | 1.5% | 0.0% | 0.0% | 21.2% | 0.0% | 0.0% | 3.5% | 0.0% | 0.0% | 0.0% | 0.0% | 2.2% | 7.1% | 0.0% | 3.1% | 3.2% | 1.6% | 0.0% | 0.1% | 0.0% | 0.0% | 0.0% | 0.000 | 0.356 | 0.435 |
| IIIB_5 | 1.2% | 0.0% | 0.0% | 11.1% | 0.0% | 0.0% | 0.1% | 0.7% | 0.0% | 0.0% | 0.0% | 1.5% | 5.8% | 0.0% | 1.6% | 2.8% | 0.0% | 0.0% | 2.1% | 0.0% | 0.0% | 0.0% | 0.020 | 0.246 | 0.270 |
| Average_IIIB | 1.7% | 0.0% | 0.0% | 16.3% | 0.0% | 0.0% | 0.9% | 0.0% | 0.0% | 3.8% | 0.0% | 0.0% | 6.2% | 0.0% | 1.7% | 4.1% | 1.2% | 0.0% | 0.7% | 0.0% | 0.4% | 0.0% | 0.020 | 0.296 | 0.370 |
| IIIT_1 | 1.9% | 0.0% | 0.0% | 26.7% | 0.0% | 0.0% | 0.6% | 1.7% | 0.0% | 0.0% | 0.0% | 4.1% | 10.7% | 0.0% | 6.2% | 6.0% | 1.3% | 0.4% | 2.7% | 0.0% | 0.0% | 0.0% | 0.000 | 0.428 | 0.624 |
| IIIT_2 | 1.1% | 0.0% | 0.1% | 30.9% | 0.0% | 0.0% | 0.3% | 0.0% | 0.3% | 0.0% | 0.0% | 0.4% | 14.6% | 0.0% | 5.8% | 2.0% | 7.4% | 0.0% | 0.0% | 1.9% | 0.0% | 0.0% | 0.000 | 0.385 | 0.648 |
| IIIT_3 | 0.0% | 0.4% | 0.0% | 20.6% | 0.0% | 0.0% | 2.3% | 0.0% | 0.3% | 0.2% | 0.0% | 0.7% | 17.3% | 0.0% | 3.1% | 2.9% | 5.8% | 0.0% | 0.4% | 0.0% | 0.0% | 0.0% | 0.000 | 0.347 | 0.539 |
| IIIT_4 | 0.0% | 0.0% | 0.0% | 10.3% | 0.0% | 0.0% | 0.5% | 0.0% | 0.0% | 0.0% | 0.0% | 0.0% | 5.3% | 0.0% | 2.1% | 1.0% | 1.6% | 0.0% | 0.2% | 0.0% | 0.0% | 0.0% | 0.000 | 0.389 | 0.211 |
| IIIT_5 | 1.4% | 0.0% | 0.0% | 25.3% | 0.0% | 0.0% | 2.0% | 1.2% | 0.0% | 0.0% | 0.0% | 0.0% | 13.2% | 0.0% | 3.7% | 6.5% | 4.3% | 0.0% | 0.0% | 0.6% | 0.0% | 0.0% | 0.000 | 0.338 | 0.583 |
| Average_IIIT | 0.6% | 0.0% | 0.0% | 26.0% | 0.0% | 0.0% | 1.7% | 0.1% | 0.2% | 0.0% | 0.0% | 0.6% | 12.4% | 0.0% | 4.3% | 4.9% | 3.7% | 0.0% | 0.0% | 0.2% | 0.0% | 0.0% | 0.000 | 0.379 | 0.548 |

**Table S4**. CIBERSORT deconvolution for each sample and immune cell type. Each value represents proportion of immune cells contributing to sample. For **Fig. 4B**, samples were consolidated for clarity of visualization. $CD4^+$ T cells include memory, activated, Tfh, Treg, and gamma delta T cells.  Average p-values for infected samples are all <0.05, p-values for saline samples are 1.

Table Legends for excel Tables.

**Table S2. 7092 differentially expressed genes >2 fold change, with padj < 0.05 for all groups compared to saline controls.** Ensembl IDs listed with $log_s$ fold values for each condition: Type II Bystanders, Type II TINs, Type III Bystanders, Type III TINs. Additional tabs list upregulated and downregulated DEGs.

**Table S3. All Canonical Pathways listed from IPA for each group compared to saline controls.** DEGs from DESeq2 were uploaded to IPA for pathway analysis with either all pathways or immune pathways. Z scores were reported if p-value < 0.05.

**Table S5. *T. gondii* transcript abundance in TINs, type II versus type III.** List of *T. gondii* transcripts that were significantly upregulated compared to saline control and then normalized between strains (type II versus type III) to show relative abundance between II-Cre TINs and III-Cre TINs. Comparison visualized in **S3**.
